# Homologous recombination defects in Shwachman-Diamond syndrome and Diamond-Blackfan anemia

**DOI:** 10.1101/2020.08.13.250068

**Authors:** Elif Asik, Nimrat Chatterjee, Alison A. Bertuch

## Abstract

Shwachman-Diamond syndrome (SDS) and Diamond-Blackfan anemia (DBA) are ribosomopathies characterized by impaired hematopoiesis and cancer predisposition. The mechanisms underlying cancer predisposition in these disorders are not well understood. We found that LCLs derived from patients with SDS or DBA had a prolonged DNA damage response and hypersensitivity to ionizing radiation, suggesting impaired DNA double strand break (DSB) repair. Consistent with this, depletion of SBDS and RPS19, the most common etiologic factors in SDS and DBA, respectively, resulted in reduced homologous recombination (HR) in HCT116 and U2OS cells. Surprisingly, depletion of EFL1, which functions with SBDS in ribosome biogenesis, did not impair HR and depletion of eIF6, which restores ribosome joining in SBDS-depleted cells, did not rescue the HR defect associated with SBDS depletion. Instead, we found SBDS and RPS19 recruitment to sites of DSBs suggesting that SBDS and RPS19 have more proximate roles in regulating HR, independent of their ribosomal functions. We propose that reduced HR shifts DSB repair toward error-prone NHEJ and this may contribute to oncogenesis in SDS and DBA. Additionally, we found SBDS and RPS19 depleted cells were hypersensitive to PARP inhibition, potentially uncovering a therapeutic target for SDS- and DBA-associated malignancies.

## INTRODUCTION

All cells rely on ribosomes for the translation of genetic information into functional proteins. The biogenesis of mammalian ribosomes, which are composed of 4 rRNAs and 80 ribosomal proteins (RPs), initiates in the nucleolus, where the pre-40S and pre-60S particles are assembled, and continues through to completion in the cytoplasm, where the exported pre-ribosomal subunits are subject to quality control steps to form the mature small 40S and large 60S subunits (1). During translation initiation, the Shwachman-Bodian-Diamond Syndrome protein (SBDS) allosterically activates elongation factor-like GTPase 1 (EFL1) to mediate the release of ribosome anti-association factor eIF6 from the 60S subunit, enabling the 40S and 60S ribosomal subunits to join to form the functional 80S ribosome and for translation to proceed (2, 3).

The ribosomopathies are a group of rare syndromes in which the production of ribosomes is perturbed due to germline pathogenic variants in genes that encode either ribosome biogenesis factors or RPs (4). One of these is Shwachman-Diamond syndrome (SDS), an autosomal recessive multisystem disorder with major clinical features of neutropenia, exocrine pancreatic dysfunction, metaphyseal dysplasia, and short stature (5). Approximately 90% of patients with SDS have pathogenic variants in *SBDS* (6), and, recently, some patients have been found to harbor pathogenic variants in *EFL1* (7–9). Consistent with the roles of SBDS and EFL1 in the eviction of eIF6 from the surface of the 60S ribosomal subunit, cells from patients bearing *SBDS* or *EFL1* mutations have defects in ribosomal subunit joining (8, 10, 11), and cells with deficient or mutant SBDS protein have a global reduction of protein synthesis (12, 13).

Another ribosomopathy that impacts the hematopoietic system is the predominantly autosomal dominant disorder Diamond-Blackfan anemia (DBA). DBA is characterized by macrocytic anemia with an absence or paucity of erythroid precursors in the bone marrow, and craniofacial, cardiac, genitourinary, and skeletal malformations (14). Approximately 80% of patients with DBA have a pathogenic variant in one of 20 genes encoding proteins of both the small 40S (RPS) and large 60S (RPL) ribosomal subunits. *RPS19* is most commonly mutated, accounting for approximately 30% of affected patients, and, among the remaining DBA-associated RP genes, only *RPL5*, *RPS26*, *RPL11*, *RPS24*, *RPS17*, *RPL35a*, *RPS15*, and *RPS10* are mutated in 1% or more of patients (15, 16). Loss-of-function variants in these genes result in reduced numbers of ribosomes, which has been shown to alter translation of a select set of transcripts in hematopoietic cells, including that of GATA1, a master transcription factor in erythropoiesis (17).

Of critical prognostic importance, patients with SDS or DBA manifest an increased risk of certain cancers throughout their lifetime. For example, patients with SDS experience myelodysplastic syndrome (MDS) and acute myeloid leukemia (AML) at a cumulative rate of 18.8% at 20 years and 36.1% by 30 years (18). For patients with DBA, the rate of MDS may be as high as 50% by 30 years (19). Patients with DBA are also prone to solid tumors, having a greater than 40-fold relative risk of colon cancer and osteosarcoma compared to the general population (19). The mechanisms underlying cancer predisposition in these disorders remain poorly defined.

In this study, we report an influence of SBDS and RPS19 on DNA double strand break (DSB) repair, and, specifically, on the high-fidelity homologous recombination (HR) repair pathway. We propose that reduced HR in SDS and DBA shifts repair of DSBs toward the error-prone nonhomologous end-joining (NHEJ) pathway, which may contribute to oncogenesis in these cancer predisposition syndromes.

## RESULTS

Our investigations were motivated by an infant who was initially evaluated in our hospital for severe chronic neutropenia, anemia, and failure to thrive. A DNA repair disorder was included in the differential diagnosis, and for this reason, the patient’s blood was sent to the Molecular Pathology Laboratory at University of California at Los Angeles for clinical radiation sensitivity testing via the colony survival assay (20, 21). An EBV-transformed lymphoblastoid cell line (LCL) was established and colony survival measured 2 weeks after exposure to 1 Gy of irradiation (IR). They reported a decreased survival fraction of 16%, placing the cells in the radiosensitive range of 14±7% (normal range 50±13%). While awaiting these results, the patient was diagnosed with SDS based on clinical features and the presence of two pathogenic, presumed biallelic *SBDS* variants (NM_016038.4: c.183_184 TA>CT and c.248+2 T>C). The combined findings drew our attention to a potential connection between SBDS and DNA repair, which was further supported by a subsequent brief report that demonstrated delayed single strand break repair after IR in LCLs derived from two patients with SDS (22).

### SDS LCLs are hypersensitive to IR and have a delayed resolution of γ-H2AX foci

To determine whether hypersensitivity to IR was a general feature of SDS LCLs, we performed colony survival assays on LCLs derived from an additional four patients with SDS, as well as an independently derived LCL from our initial patient, and a comparable number of unique healthy controls (Supplemental Table 1). Each SDS LCL was markedly sensitive to IR compared to the control LCLs (Figure 1A). The hypersensitivity to IR was associated with a sustained DNA damage response (DDR) as ascertained by the kinetics of IR-induced phosphorylated Ser139-histone H2AX (γ-H2AX) foci appearance and resolution, which reflect the detection and repair of DSBs, respectively (Figure 1, B and C). Whereas the number of γ-H2AX foci in the control LCLs increased at 1 hour and then decreased at 24 hours-post IR, the IR-induced γ-H2AX foci in the SDS LCLs were either sustained or further increased at 24 hours, suggesting a delay in the repair of IR-induced DSBs. Corroborating these findings, there was a greater increase in γ-H2AX level in the SDS LCLs at 1 and 24 hours-post IR compared to control LCLs (Figure 1D). Notably, γ-H2AX foci and protein levels were greater in the SDS LCLs compared to control LCLs even prior to IR (time 0, Figure 1, B–D). Taken together, these data suggest a defect in the repair of not only IR-induced DSBs but also endogenous DSBs in SDS LCLs.

**Figure 1.**
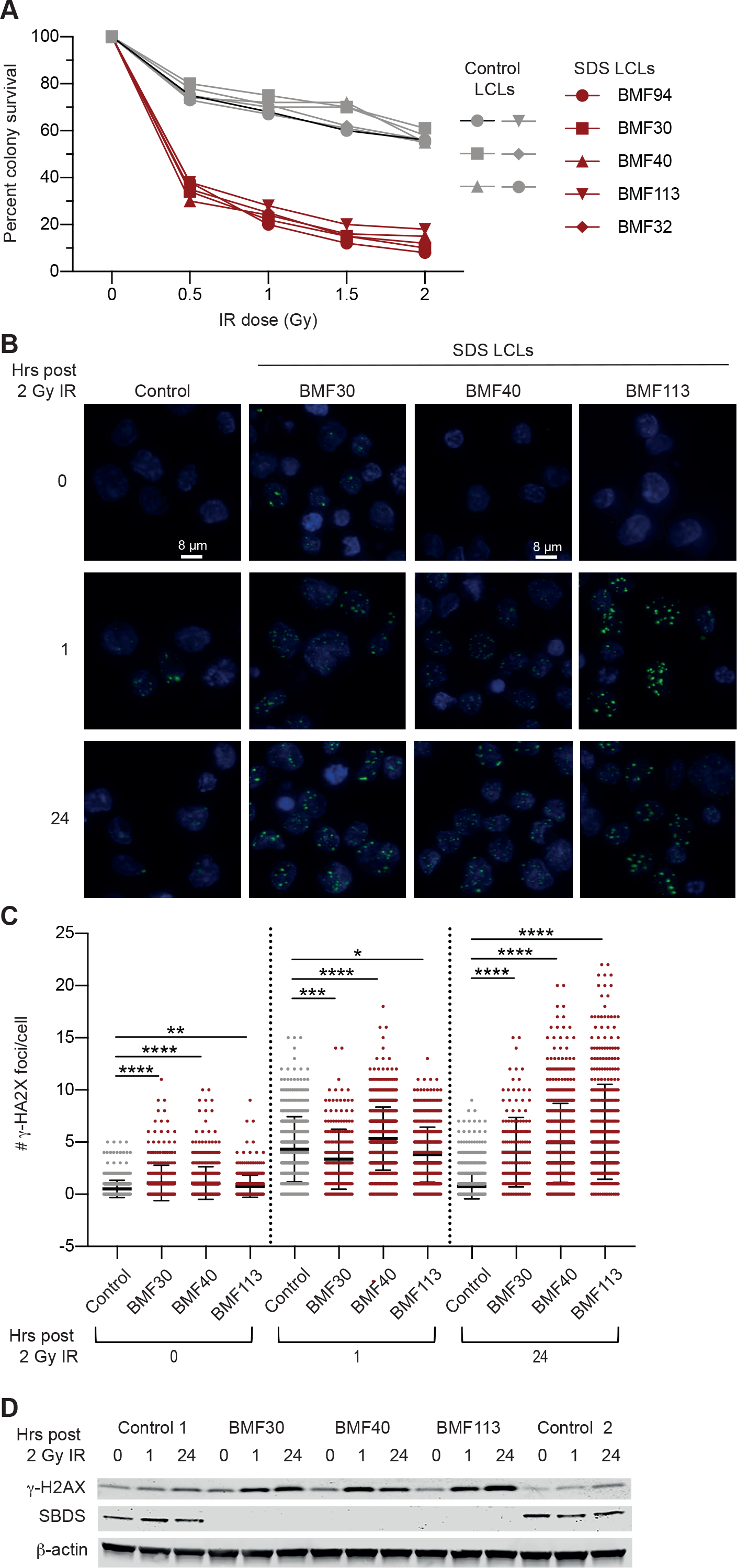
SDS LCLs are hypersensitive to IR and have a delayed resolution of γ-H2AX foci. **(A)** Colony survival assays performed with SDS and control LCLs generated from different patients with SDS (*n* = 6) or unaffected individuals (*n* = 5), respectively, pre-treatment (0) or treated with 0.5, 1, 1.5 and 2 Gy of IR. (**B)** Representative images of cells from unique SDS (BMF30, BMF40 and BMF113)- or control LCLs pre-treatment (0) and following treatment with 2 Gy IR harvested at 1 and 24 hours followed by fixation and immunolabeling with γ-H2AX antibody. DAPI was used for nuclear counterstaining. Bar, 8 μm. (**C)** Quantification of γ-H2AX foci in cells as in (B) using high throughput microscopy and single cell image analysis. Control represents the data of 3 unique control LCLs combined. Data represent mean ± standard deviation (SD) and were compared by 1-way ANOVA with Dunnett’s multiple comparisons test - \**P* < 0.05; \*\**P* < 0.01; *** *P* < 0.001; **** *P* < 0.0001. The means were calculated from 3 wells per independently grown and treated line with 200 or more cells per well. **(D)** Western blot analysis of whole cell extracts (WCE) of cells treated as in (B) probed for γ-H2AX, SBDS, and β-actin, which served as a loading control.

### SBDS-deficient HCT116 and U2OS cells show impaired HR

DSB repair by mammalian cells involves two major pathways: HR and NHEJ (23, 24). To examine these pathways, we utilized two well-characterized GFP-based reporter systems, DR-GFP and EJ5-GFP, which assess HR and NHEJ, respectively (25), in U2OS (human osteosarcoma) and HCT116 (human colon cancer) cells. Forty-eight hours after transfection with siRNAs against SBDS or a scrambled control, the cells were co-transfected with an I-Sce1 expression plasmid, to induce a DSB within the reporter, and an mCherry expression plasmid, which served as a control for transfection efficiency. Forty-eight hours later, proficiency of DSB repair was assessed by measuring the ratio of GFP-positive cells to mCherry-positive cells detected by flow cytometry. In both HCT116 and U2OS, treatment with siRNA against SBDS compared to scrambled control siRNA led to a 50–60% reduction in HR repair efficiency (Figure, 2A and B). SBDS-1 and SBDS-2 siRNAs targeted separate, unique regions of SBDS RNA, arguing against an off-target effect. In contrast, when utilizing cells bearing the NHEJ-reporter, there was no difference in NHEJ efficiency upon knockdown with SBDS-targeting siRNA compared to scrambled control siRNA (Figure 2C and D), indicating SBDS specifically impacts the HR pathway rather than having a general impact on DSB repair.

**Figure 2.**
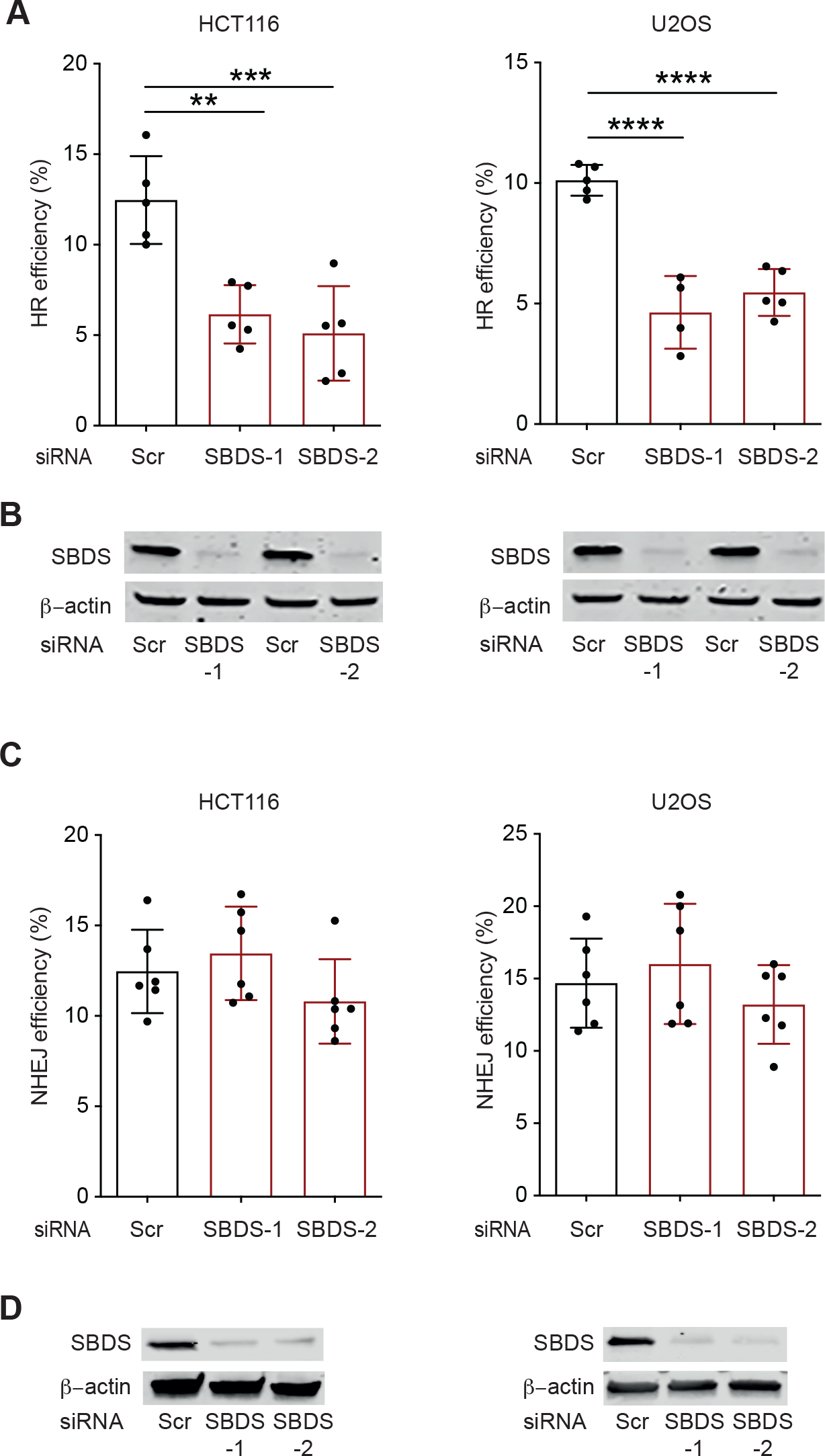
SBDS is required for proficient HR repair but not NHEJ. **(A)** HCT116 (left panel) or U2OS (right panel) cells bearing an integrated DR-GFP HR reporter (43) were transfected with a scrambled (Scr) or one of two SBDS siRNAs (−1 and −2), which targeted unique regions of SBDS RNA. After 48 hours, the cells were co-transfected with an I-Sce1-expressing plasmid to induce a DSB in the DR-GFP reporter and an mCherry-expressing plasmid as a transfection control. Forty-eight hours later, the cells were analyzed by flow cytometry. The repair efficiency was calculated by the proportion of GFP+ to mCherry+ cells. Data represent mean ± SD from 5 independent experiments compared by 1-way ANOVA and Dunnett’s multiple comparisons test - ** *P* < 0.01; *** *P* < 0.001; **** *P* < 0.0001. (**B)** Representative western blot of WCE of cells in (A) probed for SBDS and β-actin as a loading control. (**C)** The same experiment as shown in (A) except using HCT116 (left panel) or U2OS (right panel) cells bearing an integrated EJ5-GFP NHEJ reporter (25). Data represent mean ± SD from 6 independent experiments compared by 1-way ANOVA with Dunnett’s multiple comparisons test. The *P* values were not significant. (**D)** Representative western blot analysis of WCE of cells in (C).

### Defective HR in SBDS depleted cells is independent of pre-ribosomal subunit joining impairment

Having found that depletion of SBDS impaired HR, we sought to determine whether restoration of ribosomal subunit assembly in SBDS-deficient cells would restore HR. Burwick et al. demonstrated that knockdown of eIF6 improved ribosomal subunit association in the context of SBDS-deficiency (11). However, we found eIF6 depletion did not rescue the HR defect of SBDS-depleted cells (Figure, 3A and B). We considered the possibility that ribosomal dysfunction due to eIF6 deficiency might preclude rescue of HR; however, when we depleted eIF6 alone, we found that this did not impair HR. SBDS activates EFL1 to mediate the eviction of eIF6 from the 60S subunit. Therefore, as an alternative approach to address the importance of ribosome joining, we examined the impact of EFL1 depletion on HR (8). Similar to eIF6 depletion and in contrast to SBDS depletion, EFL1 depletion did not reduce HR efficiency (Figure, 3C and D). Taken together, these results suggest that the HR defect in SBDS-depleted cells is due to a novel function of SBDS distinct from its role alleviating eIF6 inhibition of ribosomal subunit joining.

**Figure 3.**
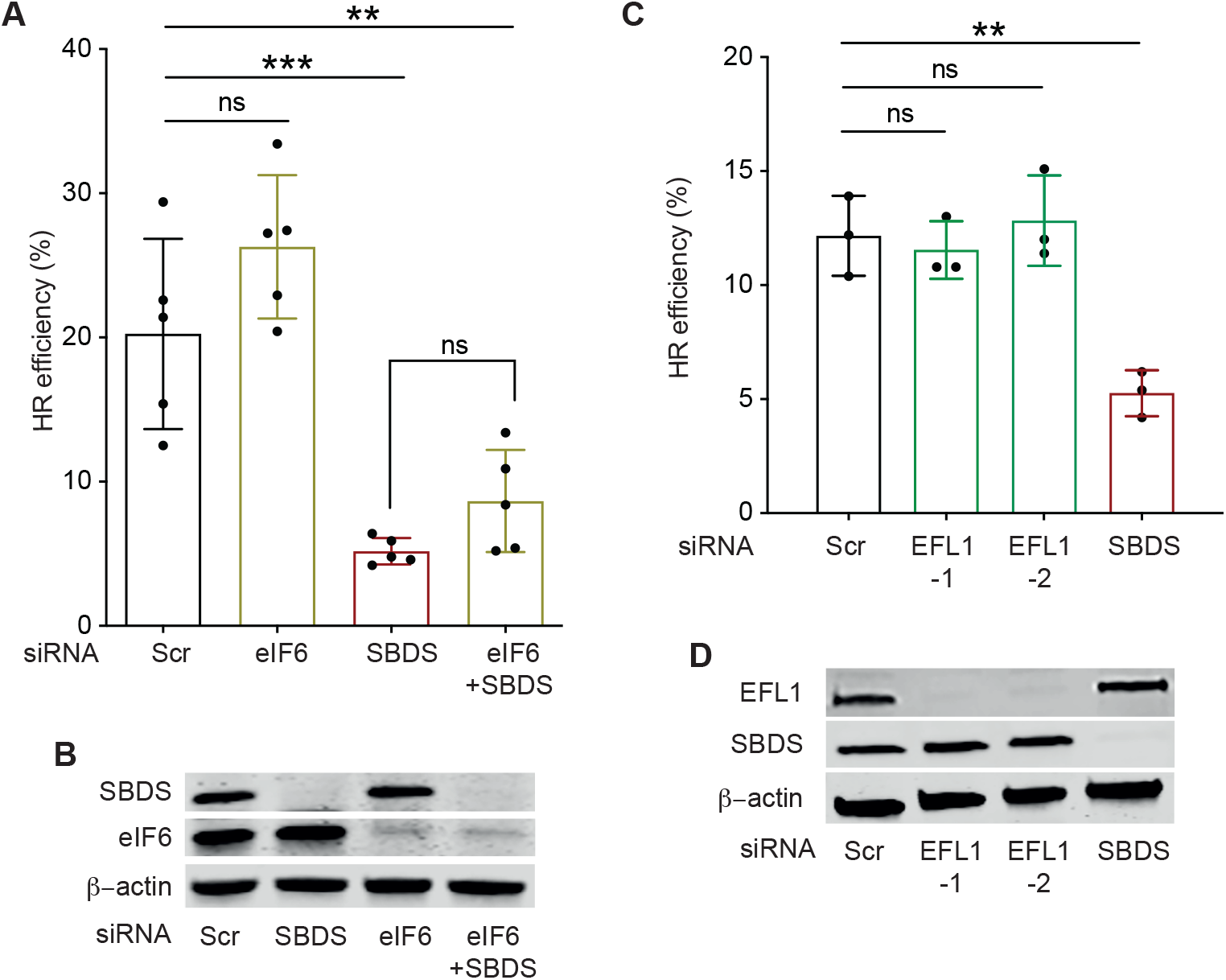
Depletion of eIF6 does not rescue and depletion of EFL1 does not phenocopy the HR defect of SBDS-depleted cells. **(A)** DR-GFP-U2OS cells were transfected with scrambled (Scr), SBDS, or eIF6 siRNAs alone or the combination of SBDS and eIF6 siRNAs and HR efficiency determined as in Figure 2A. Data represent mean ± SD of 5 independent experiments compared using a 1-way ANOVA with Tukey’s multiple comparisons test - ns *P* not significant; \*\**P* < 0.01; \*\*\**P* < 0.001. (**B)** Representative western blot of WCE of cells in (A) probed for SBDS, eIF6, and β-actin as a loading control. **(C)** As in (A) except cells were transfected with Scr siRNA, one of two siRNAs targeting distinct regions of EFL1 (−1 and −2), or SBDS. Data represent mean ± SD of 3 independent experiments compared as in (A). **(D)** Representative western blot analysis of EFL1 and SBDS protein expression of cells in (C). β-actin is loading control.

### DBA LCLs also exhibit hypersensitivity to IR and delayed resolution of γ-H2AX

To determine if IR hypersensitivity and sustained DDR is characteristic of cells with other etiologies of ribosome dysgenesis, we examined LCLs derived from patients with the ribosomal disorder DBA. We chose DBA because it is a well-established model of ribosome dysfunction due to haploinsufficiency of RPs, and patients with DBA, like those with SDS, experience hematopoietic defects and cancer predisposition (14). We utilized LCLs from patients with DBA due to different *RPS* or *RPL* gene mutations or, in one case, genetically uncharacterized DBA (Supplemental Table 1). Similar to SDS LCLs, DBA LCLs were hypersensitive to IR in colony survival assays (Figure 4A). In addition, DBA LCLs had greater numbers of γ-H2AX foci prior to IR and greater γ-H2AX foci at 24 hours following IR as compared to control lines (Figure 4, B and C). Similar findings were observed by western blot examination of γ-H2AX, which showed a greater and sustained increase in level to variable degrees in the DBA LCLs at 1 and 24 hours-post IR compared to the control LCL (Figure 4D). Thus, these data suggest a defect in the repair of endogenous and IR-induced DSBs in DBA LCLs.

**Figure 4.**
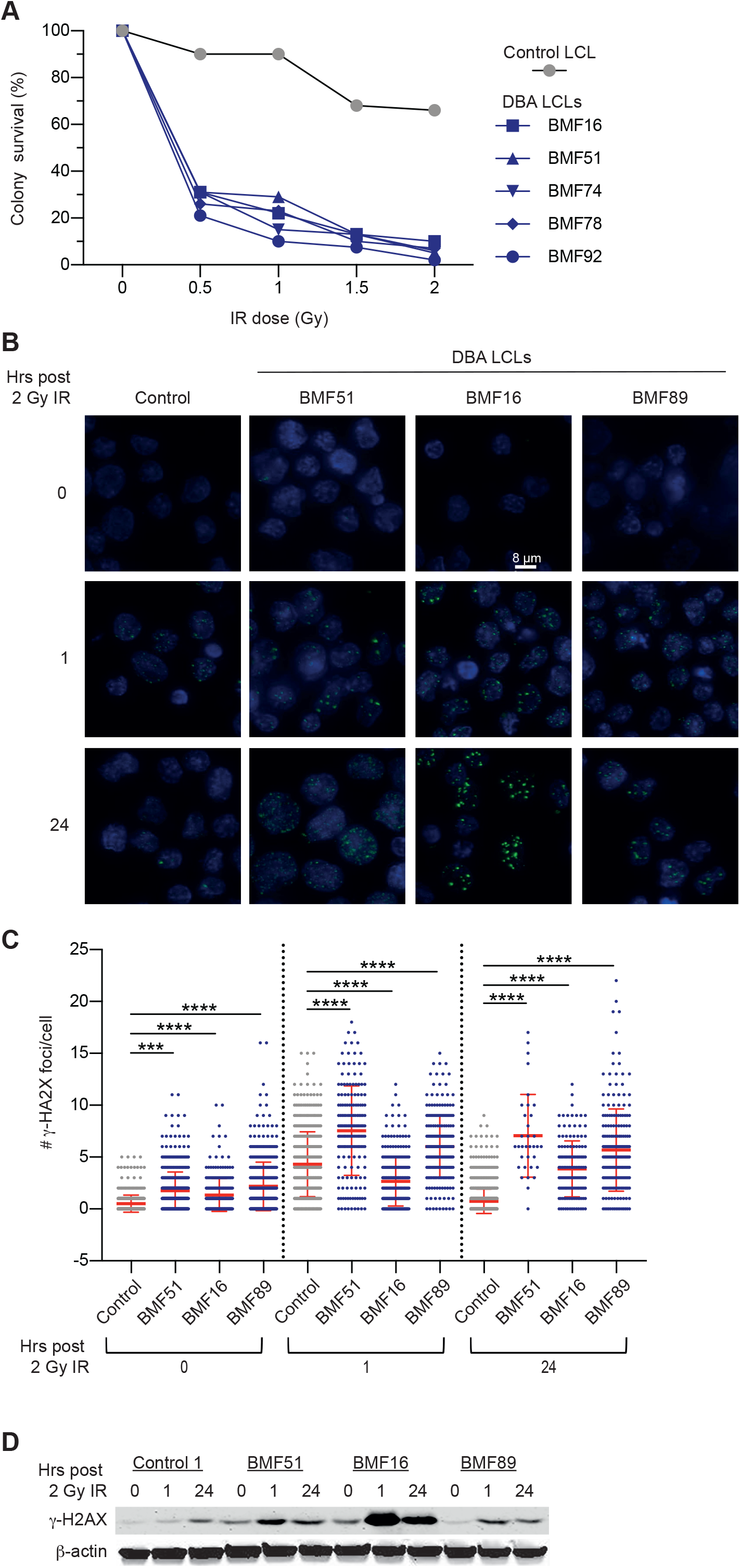
DBA LCLs also exhibit hypersensitivity to IR and delayed resolution of γ-H2AX. **(A)** Colony survival assays as in Figure 1A performed with LCLs generated from 5 patients with DBA (BMF16, BMF 51, BMF 74, BMF 78, and BMF 92, see Supplemental Table 1 for genotypes) or an unaffected control. **(B)** Representative images of cells from DBA- or a control LCL pre-treatment (0) and following treatment with 2 Gy IR and harvested at 1 and 24 hours followed analysis of γ-H2AX foci as in Figure 1B. **(C)** Quantification of γ-H2AX foci in cells treated as in (B) using high throughput microscopy and single cell image analysis. The data were acquired along with the SDS LCLs in Figure 1C and, therefore, the data for the combined controls are the same. Data represent mean ± SD compared using 1-way ANOVA with Dunnett’s multiple comparisons test - \*\*\**P* < 0.001; \*\*\*\**P* < 0.0001. **(D)** Western blot analysis of whole cell extracts (WCE) of a control and 3 DBA-LCLs treated as in (B) probed for γ-H2AX and β-actin as a loading control.

As with the SDS LCLs, we sought to determine if these findings were due to a defect in DSB repair. We chose to examine the impact of RPS19 depletion on repair of the HR and NHEJ reporters, as *RPS19* is the most commonly mutated gene in DBA. With striking similarity to the SBDS knockdown, we found a reduction in HR efficiency and preservation of NHEJ efficiency following two different siRNA-mediated knockdowns of RPS19 (Figure, 5A–D), suggesting that RPS19, like SBDS, impacts the HR pathway.

**Figure 5.**
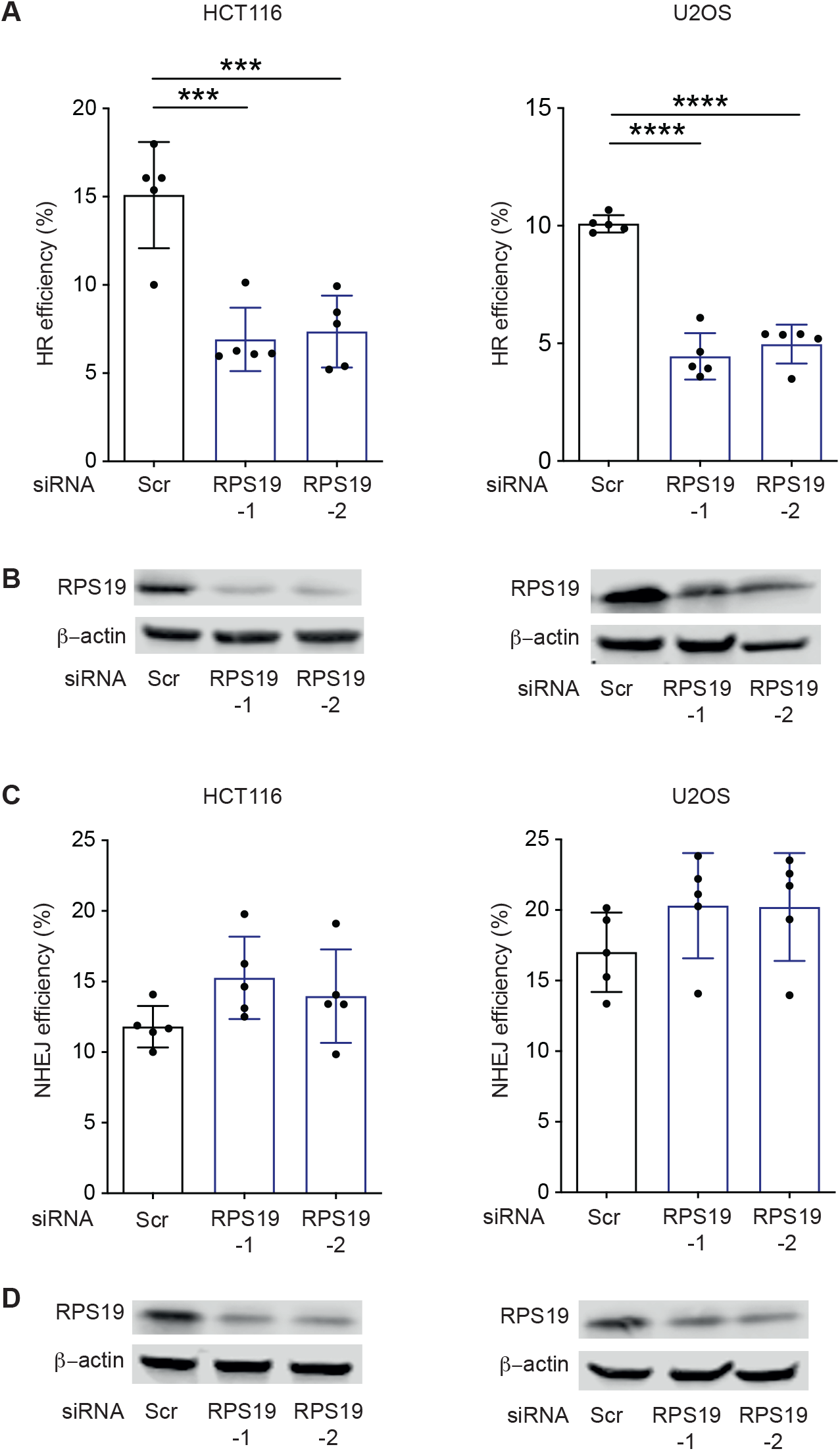
RPS19 is required for proficient HR repair but not NHEJ. **(A)** HR assays as described in Figure 2A with DR-GFP-HCT116 (left panel) or DR-GFP-U2OS (right panel) HR reporter cells except the cells were transfected with scrambled (Scr) or one of two RPS19 siRNAs targeting unique regions of RPS19 RNA (−1 and −2). Data represent mean ± SD of 5 independent experiments compared using 1-way ANOVA and Dunnett’s multiple comparisons test - \*\*\**P* < 0.001; \*\*\*\**P* < 0.0001. (**B)** Representative western blot of WCE of cells in (A) probed for RPS19 and β-actin as a loading control. **(C)** Assays as in (A) except using EJ5-HCT116 (left panel) or EJ5-U2OS (right panels) NHEJ reporter cells. Data represent mean ± SD of 5 independent experiments compared as in (A). *P* values were not significant. **(D)** Representative western blot of WCE of cells in (C) probed for RPS19 and β-actin.

### Disturbance in cell cycle progression does not account for the HR defect in SBDS- or RPS19-depleted cells

HR occurs during late S and G2 phases of the cell cycle when a sister chromatid is available. Thus, we examined whether a disruption in cell cycle progression upon SBDS or RPS19 depletion might account for the impaired HR. U2OS cells treated with SBDS, RPS19 or scrambled siRNA were collected after 72 hours for cell cycle analysis based on BrdU incorporation and 7-aminoactinomycin D (7-AAD) staining. While percentage of cells in S phase was reduced in the SBDS-depleted cells, there was increase in the percentage of G2/M cells and there was no decrease in the combined late S and G2/M percentage compared to the control (Figure 6). This result suggests that a reduction of late S/G2 cells is not a contributing factor to the decreased HR proficiency associated with SBDS depletion. The RPS19-depleted cells showed increased in percentages of cells in early and late S phase and G2/M phase compared to scramble siRNA-treated cells (Figure 6). While the effects are notable, they suggest that RPS19-dependent perturbations in HR are also cell cycle-independent.

**Figure 6.**
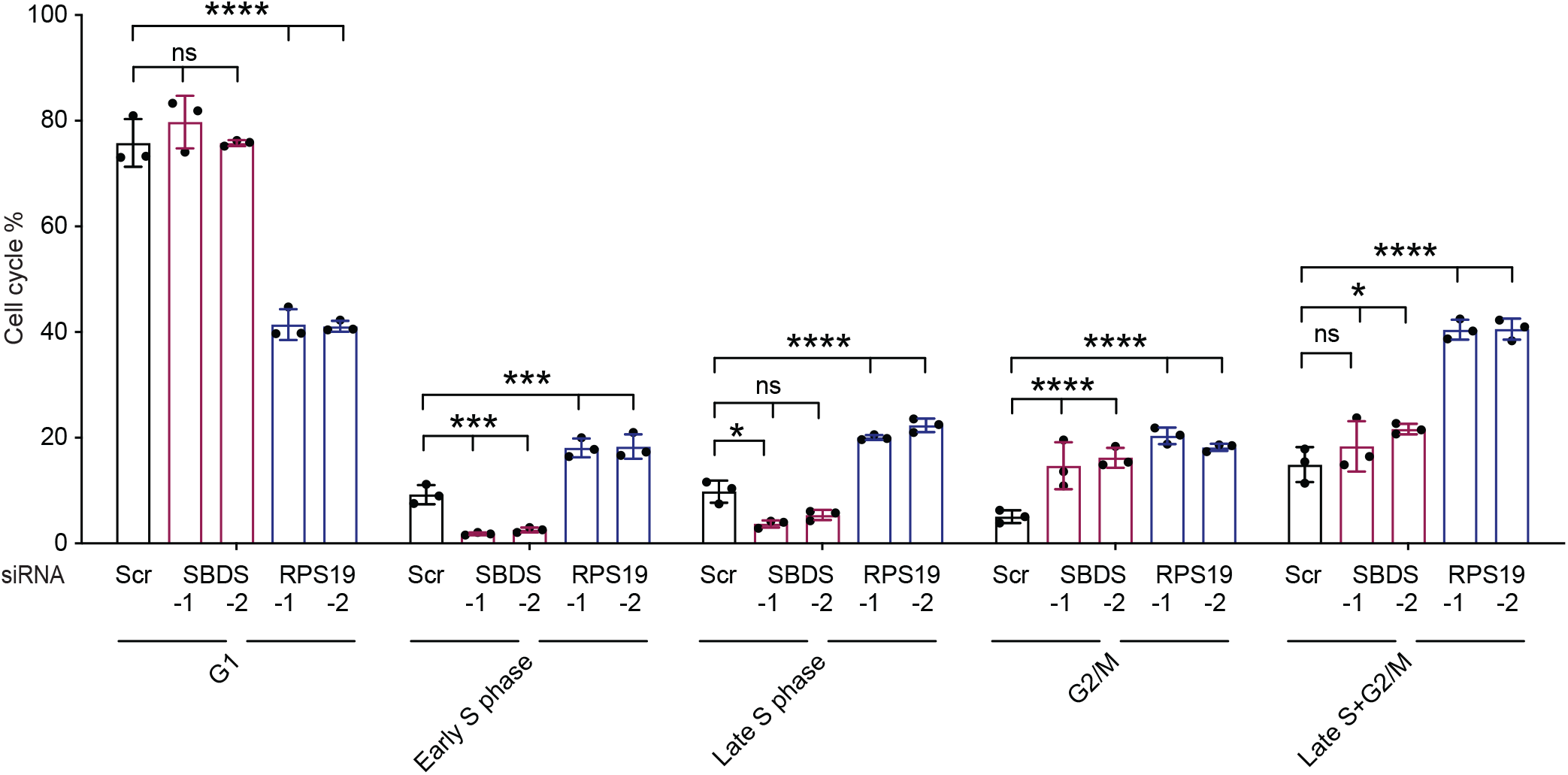
The HR defect in SBDS- or RPS19-depleted cells is not accounted for by altered cell cycle progression. U2OS cells were transfected with scrambled (Scr), SBDS or RPS19 siRNA for 72 hours, followed by flow cytometry using 7-AAD and BrdU. Data represent mean ± SD for 3 independent experiments compared using 2-way ANOVA and Dunnett’s multiple comparisons - ns *P* not significant; \*\**P* < 0.01; \*\*\**P* < 0.001; \*\*\*\**P* < 0.0001.

### Decreased RAD51 levels in SBDS- or RPS19-deficient cells

As an alternative explanation for the HR reduction in SBDS- and RPS19-deficient cells, we asked if SBDS or RPS19 deficiency resulted in a reduction of HR-specific factors. We found a reduced level of the RAD51 recombinase in SDS LCLs compared to controls both prior to and following IR (Figure 7A). Corroborating this finding, we found a reduced level of RAD51 in U2OS cells treated with SBDS siRNA compared to scrambled siRNA-treated cells (Figure 7, B and C). In contrast, a survey of several key NHEJ factors revealed no consistent reductions in the SDS LCLs (Supplemental Figure 1). A reduction in RAD51 was also observed in cells treated with RPS19 siRNA (Figure 7, D and E), once again revealing commonality between the impact of SBDS- and RPS19-depletion on HR. Importantly, the RAD51 protein level remained was unchanged in EFL1 deficient cells (Figure 7, F and G), consistent with a lack of effect of EFL1-depletion on HR (Figure 3B). We also observed that the expression of *RAD51* mRNA, measured by reverse transcriptase-quantitative PCR (qPCR), was downregulated in SBDS siRNA-treated U2OS cells compared to scrambled siRNA-treated controls (Figure 7H), suggesting the reduction in RAD51 protein may not involve impaired translation as would be expected given the ribosome biogenesis defect.

**Figure 7.**
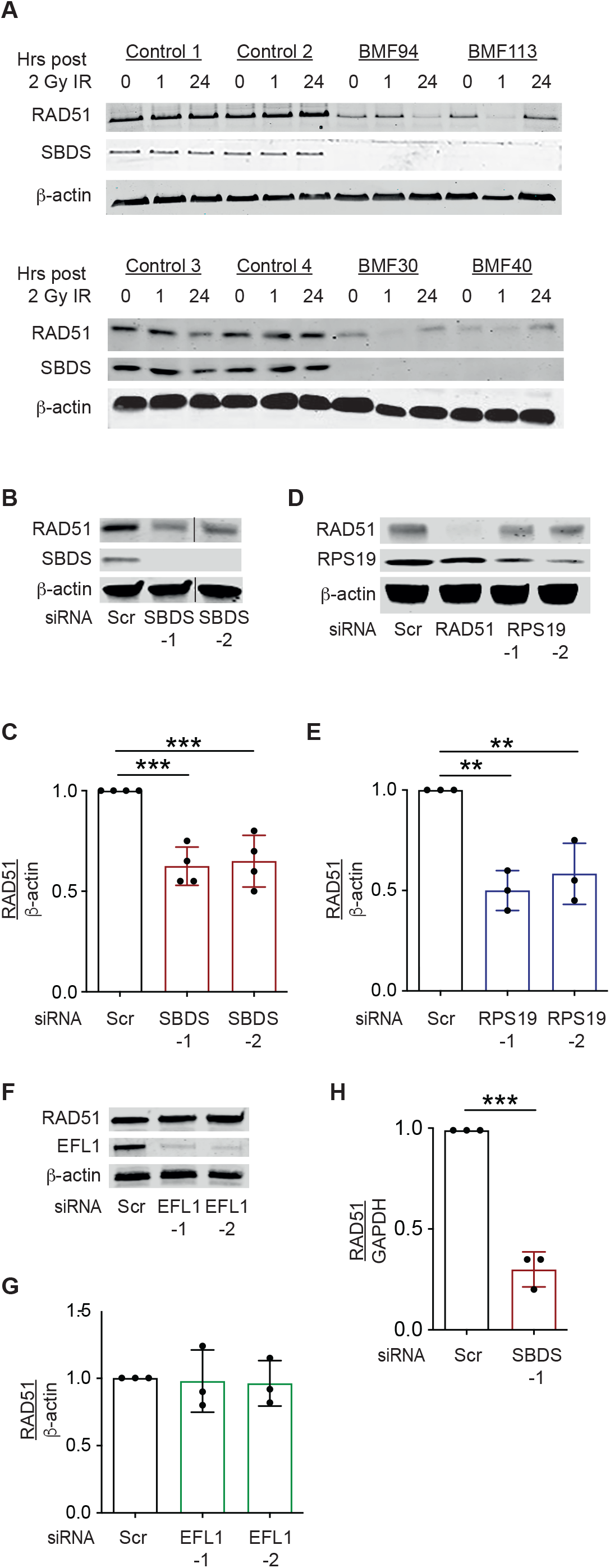
SBDS- or RSP19-deficient cells have decreased RAD51. **(A)** Representative western blot of WCEs from 4 SDS (BMF94, BMF113, BMF30 and BMF40) and 4 unique control LCLs prepared pre-treatment (0) and following treatment with 2 Gy IR harvested at 1 and 24 hours analyzed for RAD51 and SBDS proteins. β-actin is a loading control. **(B)** Representative western blot of U2OS cells transfected with Scr, SBDS-1 or SBDS-2 siRNA for 72 hours analyzed for RAD51 and SBDS proteins. β-actin is a loading control. The lines indicate the removal of a non-relevant lane from the image. **(C)** Quantification of western blot RAD51 protein measurement normalized to β-actin. Data represent mean ± SD for 4 independent experiments compared using 1-way ANOVA and Dunnett’s multiple comparisons test - \*\*\**P* < 0.001. **(D)** Representative western blot of U2OS cells transfected with Scr, RAD51, RPS19-1 or RPS19-2 siRNA for 72 hours analyzed for RAD51 and RPS19 proteins. β-actin is a loading control. **(E)** Quantification and analysis of western blots from 3 independent experiments as in B except probed for RPS19 along with RAD51 and β-actin. \*\**P* < 0.01 **(F)** Representative western blot of WCEs prepared from cells transfected with Scr, EFL-1 or EFL-2 siRNAs as in (B) and (D). **(G)** Quantification and analysis of western blots from 3 independent experiments as in (C) and (E). Data represent mean ± SD compared using 1-way ANOVA and Dunnett’s multiple comparisons test. *P* values were not significant. **(H)** U2OS cells were transfected with Scr, SBDS-1, or SBDS-2 siRNAs and 72 hours later, total RNA isolated and reverse transcribed using oligodT primer. RAD51 gene expression relative to GAPDH was determined by qRT-PCR in 3 independent experiments. Data represent mean ± SD for 3 independent experiments compared using unpaired t test - \*\*\*\**P* < 0.001.

### Restoration of RAD51 levels cells is insufficient to rescue the HR defect in SBDS-or RPS19-deficient cells

Given the impact of SBDS- and RPS19-depletion on RAD51 levels (Figure, 7A–E) and the requirement of RAD51 for HR (26), we tested whether restoration of RAD51 level could rescue the HR phenotype in SBDS- or RPS19-deficient cells. To do this, we performed I-Sce1-based HR reporter assays with forced expression of RAD51 from an exogenous plasmid. As expected, SBDS and RAD51 protein expression was reduced in U2OS cells treated with siRNA against SBDS, with RAD51 levels restored by co-transfection of an exogenous RAD51 expression plasmid (Figure 8A) Nonetheless, the HR efficiency remained reduced despite RAD51 restoration (Figure 8B). Similar results were observed in RPS19-depleted cells (Figure 8, C and D). These data indicate that an additional mechanism plays a role in the reduced HR efficiency observed in SBDS- and RPS19-deficient cells.

**Figure 8.**
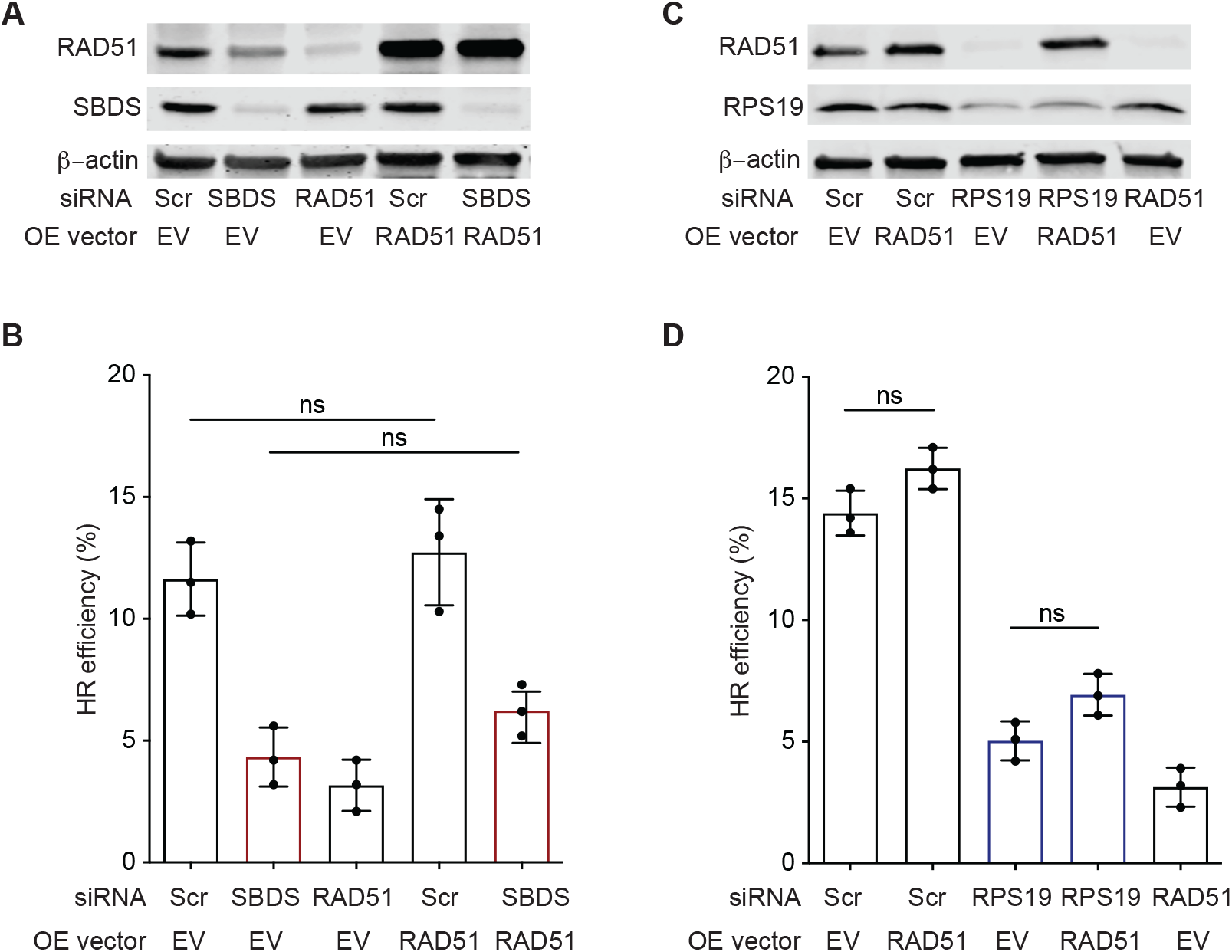
Overexpression of RAD51 in SBDS- or RPS19-depleted cells does not rescue the HR-deficiency. U2OS cells were transfected with Scr, SBDS or RAD51 siRNA. Twenty-four hours later, the cells were transfected via electroporation with an I-Sce1 expression plasmid and either an RAD51 overexpression plasmid or empty vector, both of which expressed mCherry. Forty-eight hours later, the cells were subjected **(A)** western blot analysis for RAD51, SBDS and β-actin proteins and **(B)** flow cytometry analysis for HR efficiency based on the ratio of GFP+ to RFP+ cells. Data represent mean ± SD for 3 experiments compared using 1-way ANOVA with Dunnett’s multiple comparisons test - ns, nonsignificant *P* value. **(C-D)** The same experimental approach as shown in panels (A) and (B) except that RPS19 was depleted by siRNA and analyzed by western blot. Data similarly represent mean ± SD for 3 experiments.

### SBDS and RPS19 are found in the vicinity of DSB

A recent study reported that RPL6 was rapidly recruited to sites of DNA damage induced by laser microirradiation (27). Thus, we tested whether the SBDS and RPS19 proteins might also function more directly in DSB repair through localization to DSBs. To test this hypothesis, we utilized U2OS cells containing an integrated DR-GFP transgene, as for the HR assays (Figure 2A) and monitored SBDS and RPS19 recruitment following I-Sce1-DSB induction via ChIP followed by qPCR. ChIP-qPCR analysis revealed significant association of SBDS surrounding the induced DSB (Figure 9A). Analysis of RPS19 also showed association, albeit with a different enrichment pattern. (Figure 9B). These data suggest that SBDS and RPS19 have a more proximate role in the response to DSBs.

**Figure 9.**
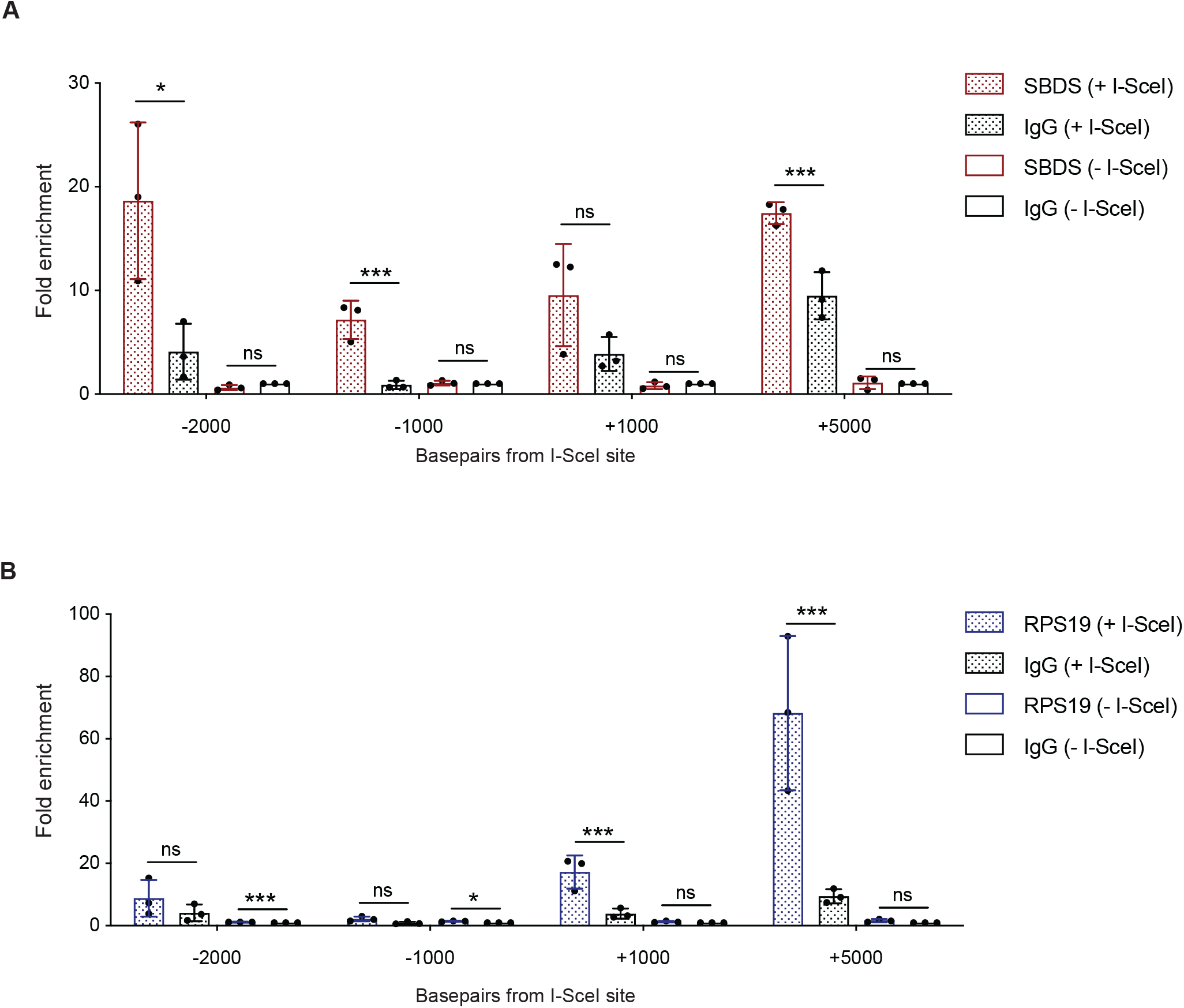
SBDS and RPS19 are found in the vicinity of DSB. **(A)** DR-GFP-U2OS cells were transfected with either an I-Sce1 expression plasmid or an empty vector. ChIP was performed 24 hours later with an SBDS antibody. IgG was used as negative control. The DNA was analyzed by qPCR using primers designed for the designated regions on the DR-GFP vector and the fold of enrichment of the corresponding region relative to GAPDH was calculated and normalized to the value obtained for IgG (-I-Sce1) of the same region. Data represent mean ± SD for 3 independent experiments using t tests comparing the normalized fold enrichment of SBDS (+I-Sce1) to IgG (+I-Sce1) and SBDS (-I-Sce1) to IgG (-I-Sce1) for each region. \**P* < 0.05; \*\**P* < 0.01; \*\*\**P* < 0.001; \*\*\*\**P* < 0.0001. **(B)** Same as (A) except that ChIP was performed with an RPS19 antibody.

### SBDS- and RPS19-depleted cells are hypersensitive to poly (ADP-ribose) polymerase (PARP) inhibition

Cells deficient in HR are sensitive to inhibition of PARP, a vulnerability which has been exploited in cancer therapeutics (28, 29). We therefore examined whether SBDS- or RPS19-deficient cells were sensitive to olaparib, a PARP inhibitor used in the treatment of *BRCA-* mutated cancers. Similar to RAD51-depleted cells, SBDS- and RPS19-depleted cells had decreased viability after 48 hours of olaparib treatment in a dose-dependent fashion compared to the scrambled siRNA treated control cells (Figure 10). These results corroborate a HR deficiency resulting from SBDS- and RPS19-deficiency.

**Figure 10.**
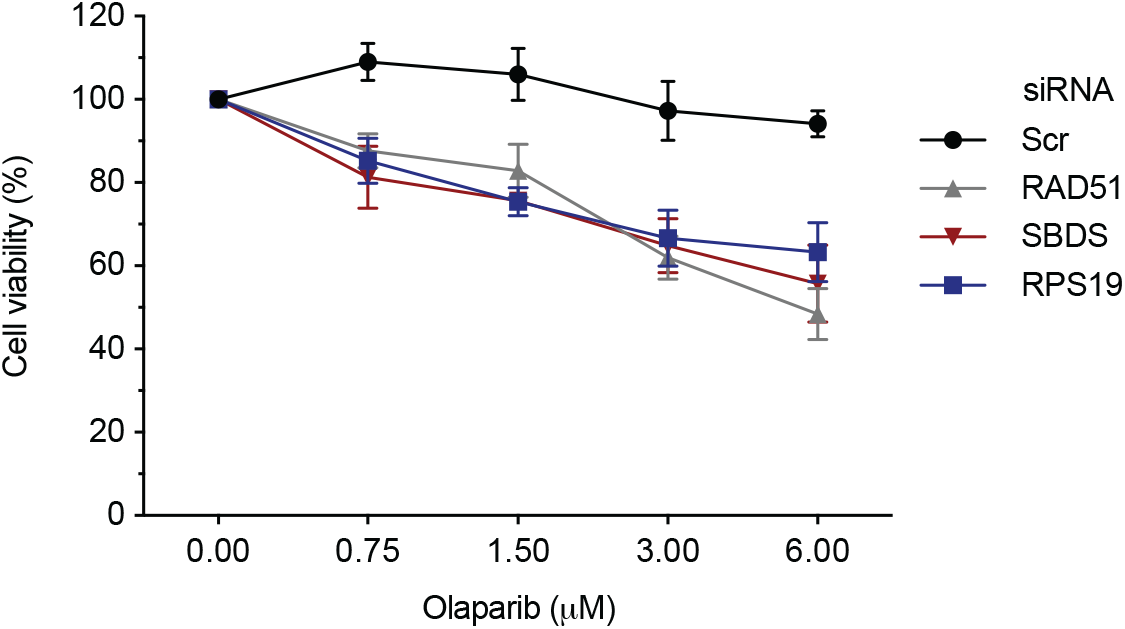
SBDS- and RPS19-depleted cells are hypersensitive to poly (ADP-ribose) polymerase (PARP) inhibition. U2OS cells were transfected with Scr, SBDS, RAD51 RPS19 or siRNA. After 24 hours, the cells were replated and further cultured in the presence of varying concentrations of olaparib (0.75, 1.5, 3 and 6 μM) or the DMSO diluent alone (0 μM). Viability was assessed 48 hours later. Data represent mean ± SD of 3 independent experiments with 3 technical replicates.

## DISCUSSION

Many studies have sought insight into the function of the SBDS protein, with most focused on its role in ribosomal subunit joining (reviewed in 30). A smaller number have described its impact on processes apparently unrelated to ribosome biogenesis, such as mitotic spindle stabilization, neutrophil chemotaxis, regulation of the production of reactive oxygen species and mitochondrial function, cellular stress responses, and telomerase recruitment (13, 31–34). Similarly, some RPs have been implicated in processes independent of translation, most notably regulation of p53 degradation (27). Here, we demonstrate that SBDS and RPS19 also have roles in DNA DSB repair. We found LCLs derived from patients with SDS due to *SBDS* mutation or patients with DBA due to a variety of RP gene mutations, including *RPS19*, have an impaired DDR and are hypersensitive to IR (Figure 1 and 4). Corroborating and extending these findings, we found SBDS- and RPS19-deficient cell lines are specifically impaired for the HR pathway of DSB repair (Figure 2 and 5). Given the importance of HR in the maintenance of genome stability and tumor suppression (35), these findings raise the possibility that impaired DSB repair contributes to the hematopoietic defects or cancer predisposition in SDS and DBA.

How might SBDS and RPS19 influence HR? We found levels of RAD51 protein were diminished in SBDS- and RPS19-depleted cell lines, but exogenous expression of RAD51 failed to rescue the HR deficiency in these cells (Figure 8). Thus, additional factors must be contributing to the HR defect. Given the impact of SBDS and RPS19 on global translation, it is possible that translation of other HR factors is impaired upon SBDS or RPS19 knockdown, thereby reducing HR. However, several lines of evidence in this study suggest that these proteins may influence HR independently of their impact on translation. Specifically, eIF6 depletion did not rescue the HR defect, as would be expected if ribosome subunit joining was restored (Figure 3A); EFL1 depletion did not phenocopy the HR defect of SBDS-deficient cells (Figure 3B), as would be expected if impaired ribosome subunit joining was central to the phenotype; and both SBDS and RPS19 were recruited to sites of an induced DSB (Figure 9). While the association of these proteins with DSBs is novel, a few other ribosomal proteins, RPL6, RPL8 and RPS14, but not RPL9, have been shown previously to be recruited to sites of DNA damage-induced by laser microirradiation (27). RPL6 recruitment occurs via interaction with H2A and in a PARP-dependent fashion. Further studies are needed to elucidate factors that govern SBDS and RPS19 recruitment, as well as studies to determine the impact on events downstream of DSB-induction and γ-H2AX phosphorylation. Notably, in contrast to SBDS- and RPS19-depletion, RPL6-depletion reduces both HR and NHEJ, thus, differences are likely to be uncovered.

The mechanism(s) underlying cancer predisposition in SDS and DBA are of fundamental biological and clinical interest. A prominent model for the increased incidence of leukemia in SDS relates to p53 tumor suppressor function. MDM2 is an E3 ubiquitin-protein ligase that ubiquitinates p53, leading to its degradation by the proteasome. Under conditions of ribosome biogenesis stress, the 5S RNP (composed of RPL5, RPL11, and 5S rRNA), or other free RPs bind to MDM2 and competitively inhibit the binding of p53, thereby alleviating MDM2-induced p53 degradation (36). This activation of p53 secondary to ribosome biogenesis stress has been proposed to occur in SDS, restricting proliferation and inducing apoptosis of hematopoietic stem cells (37). Subsequent acquisition of a *TP53* mutation would attenuate the restriction, and a somatic mutation in the other *TP53* allele would further drive genomic instability and leukemogenesis. Consistent with this model, p53 protein is stabilized in the bone marrow of patients with SDS (38), clonal hematopoiesis due to *TP53* mutations is observed in approximately 50% of patients with SDS (39), and *TP53* loss-of-function mutations have been universally observed in SDS-associated MDS/AML (40) (although the numbers sequenced remain small to date).

Our findings raise the possibility that impaired HR, with secondary accumulation of mutations, may also play a role in cancer predisposition. In addition to showing persistent γ-H2AX foci following IR, we found an increase in γ-H2AX foci and protein levels in untreated SDS and DBA cells, presumably as a result of endogenous DSBs (Figure 1 and 4). Because an intact DDR is required during cell proliferation and tissue maintenance, its persistent impairment would result in a slow accumulation of DNA damage. HR is the most faithful pathway of DSB repair because it uses the homologous sister chromatid as a template (41). The NHEJ pathway, which does not rely on a DNA template and frequently requires processing of the end structures to permit end ligation, is more error-prone. Thus, even in the absence of exogenous damage, a shift in repair from HR to NHEJ may result in a mutagenic phenotype.

As noted above, we found that the depletion of EFL1 did not result in an HR defect (Figure 3B). This would suggest that, if the HR defect associated with *SBDS* mutation contributes to malignant progression, then those with SDS due to *EFL1* mutations would not have an increased risk of MDS/AML. None of the 10 patients with biallelic *EFL1* variants and SDS have been reported to have MDS or AML (7–9). Nevertheless, the numbers of patients are too small to draw conclusions at this time.

Patients with SDS who develop MDS or AML have poor outcomes, due in part to therapy resistance (42). As would be expected for cells with impaired HR, we found that SBDS- and RPS19-depleted cells had increased sensitivity to PARP inhibition. This raises a possible role for PARP1 inhibitors in the treatment of cancers in patients with SDS and DBA.

## METHODS

### Cell lines and culture and plasmids

LCLs were previously generated under Baylor College of Medicine IRB-approved study H-17698 using the tissue culture core laboratory within the Department of Molecular and Human Genetics, Baylor College of Medicine. LCLs were cultured in RPMI 1640 medium containing L-glutamine (Invitrogen) and 10% FBS in 5% CO_2_. HCT116 and U2OS cells (ATCC) were maintained in McCoy’s medium supplemented with 10% FBS.

DR-GFP-HCT116 and DR-GFP-U2OS reporter cells were generated by electroporating HCT116 and U2OS cells with Sac1/Kpn1 linearized pHPRT-DRGFP (43) (Addgene, plasmid #26476) DNA, respectively. Puromycin-resistant clones were pooled 10–15 days later and used for HR assays. EJ5-GFP HCT116 cells and EJ5-GFP U2OS reporter cells were similarly generated by selecting stable clones after electroporation of HCT116 and U2OS cells with XhoI linearized pimEJ5-GFP (25) (Addgene, plasmid 44026) DNA, respectively.

The RAD51 overexpression plasmid (pAB1118) was constructed by cloning *RAD51* cDNA into the pCAGGS-mCherry plasmid (Addgene, plasmid #41583). First, a 2.4 kb NotI/BamHI *RAD51* cDNA-containing fragment from pENTR221 (Invitrogen) was subcloned into pEF6/HisA (Invitrogen), placing *RAD51* under the control of an EF1α promoter to created pAB1116. The EF1α promoter-*RAD51* cDNA unit was PCR amplified and subcloned into the SfiI site of pCAGGS-mCherry to generated pAB1118.

### Colony survival assays

Colony survival assays were performed as previously described (21).

### Immunofluorescence assays

LCLs were grown to >75% confluence and irradiated with a single dose of 0 and 2 Gy on 6 well dishes using a Rad Source Pro Biological Irradiator Model 2000 (X-ray, 25 mA, 160 kV [peak]). Cells were harvested 1 and 24 hours after IR, fixed in 4% paraformaldehyde for 20 minutes on ice, quenched with ammonium chloride 0.1 M in PBS for 15 minutes at room temperature (RT) and spotted in 384-well glass bottom plates by cytospin at 1000 rpm for 3 minutes. Cells were then permeabilized with PBS/Triton-X100 0.1% for 20 minutes at RT. After 30 minutes of blocking in 5% milk/TBST, cells were incubated at 4 °C with anti-phospho-histone H2A.X (Ser139) monoclonal mouse antibody (Millipore, clone JBW301) at 1:1000 dilution. After washing, cells were incubated at RT with Alexa488-conjugated anti-mouse IgG antibody (Life Technologies, A28175) at 1:1,000 dilution for 30 minutes and counterstained with DAPI (2 μg/mL). The number of γ-H2AX foci per nucleus was measured from cells imaged with a GE Healthcare DeltaVision deconvolution microscope, using an Olympus 40x/0.95NA PlanApo objective. Z-stacks (0.35 μm steps) were acquired, covering the entire nucleus, and projected by maximum intensity after restorative deconvolution. First, nuclear masks were defined using a watershed algorithm based on the DAPI signal. Foci were identified using a local maxima detection algorithm after denoising and gaussian filtering the γ-H2AX. The number of foci within each nuclear mask was then recorded. This analysis was performed with MATLAB version 2016b.

### Western blotting

Cells were lysed in RIPA buffer (50 mM Tris-HCl pH 8.0, 150 mM NaCl, 1% Triton X-100, 0.5% Na deoxycholate, and 0.1% SDS) with 1x protease inhibitor cocktail III (Calbiochem) for 30 minutes on ice, sonicated on ice for 5 minutes and then centrifuged at 14,000 rpm for 15 minutes at 4°C. The protein concentration of the supernatant was determined using the bicinchoninic acid (BCA) protein assay kit (Pierce). Thirty to 40 μg of protein were loaded per sample. The immunoprecipitates were resolved on 4-12% Criterion XT Bis-Tris or 4-20% Mini-Protean TGX precast gels (BioRAD) and transferred to Immobilon-FL PVDF membranes (Millipore). The following antibodies were used for western blotting and immunoprecipitation: polyclonal rabbit anti-SBDS (Abcam, ab128946), polyclonal rabbit anti-RPS19 (Bethyl-A304-002A), rabbit anti-EFTUD1 (Bethyl A305-854A-T), rabbit anti-RAD51 (Abcam ab176458), polyclonal rabbit anti-eIF6 (Bethyl A303-029A-M), monoclonal mouse anti-phospho-Histone H2A.X (Ser139) antibody (Millipore, clone JBW301) and monoclonal mouse anti-β-actin (Sigma-Aldrich, A5441). Protein band quantification was performed using a LI-COR Odyssey CLx Infrared Imaging System and Image studio lite software.

### HR and NHEJ repair reporter assays

DR-U2OS and -HCT116 cells and EJ-U2OS and -HCT116 reporter cells were plated in 6 well plates and transfected with 5 μL Lipofectamine RNAiMAX transfection reagent (Thermo Fisher) and siRNAs (Invitrogen) (sequences provided in Supplemental Table 2) at 25 nM final concentrations according to the manufacturer’s instructions. Forty-eight hours later, the cells were co-electroporated with 10 μg pCBA-SceI (Addgene, plasmid #26477) (44) and 1 μg pCAGGS-mCherry (Addgene, plasmid #41583) (45). The cells were harvested 2 days later and subjected to flow cytometry analysis to detect GFP-positive and mCherry-positive cells using a BD LSRII flow cytometer and BD FlowJo software. The repair efficiency was scored as the percentage of GFP- vs mCherry-positive cells.

### Cell cycle analysis

Cells were grown for 1 hour in medium supplemented with 10 μM BrdU. Cells were fixed with 70% ethanol and incubated at −20°C for at least 2 hours. The DNA was denatured using 1 mL of 2N HCl/Triton X-100 and then neutralized with 500 μL 0.1M Na_2_B_4_O_7_. Cells were blocked using PBS/1% BSA/0.5% Tween-20 and incubated with monoclonal mouse FITC anti BrdU (Biolegend, clone 3D4) at 1:50 dilution for 20 minutes. Cells were suspended in 1 mL of PBS containing 1μg/mL 7-ADD viability staining solution, incubated at RT for 30 minutes, and then analyzed by FACS on a BD LSRII flow cytometer and analyzed using FlowJo software.

### RNA analyses

RNA was isolated from U2OS cells using the RNeasy® kit (QIAGEN) and cDNA synthesized using the qScript Flex cDNA Synthesis Kit (Quanta Biosciences). The following primer pairs were used: RAD51 [forward (for)] 5’GGCAATGCAGATGCAGCTTGAAGC3’ and [reverse (rev) 5’CCGTGAAATGGGTTGTGGGCCA and GAPDH 5’CACATGGCCTCCAAGGAGTAAG and 5’TACATGACAAGGTGCGGCTCCC. Quantitative PCR was performed on a QuantFlex 6 system (Applied Biosystems) using PowerUp SYBR Green Master Mix (Applied Biosystems) according to the manufacturer’s protocol. The relative changes in RNA abundance were calculated by comparative ΔCt method with normalization to GAPDH.

### ChIP assays

Briefly, DR-U2OS cells were electroporated with 10 μg pCBA-SceI or pCAGGS-mCherry as a vector control. After 24 hours, the cells were fixed with formaldehyde (final concentration 1% vol/vol) for 10 minutes, followed by the addition of glycine (final concentration of 125 mM) to stop the reaction. Cells were washed three times with ice-cold PBS, then scraped from culture dishes into a microfuge tube. Cells were collected by centrifugation at 5000 × g for 10 minutes at 4°C, lysed with RIPA lysis buffer with protease inhibitors on ice for 15 minutes and then sonicated with a Bioruptor (Diagenode) for 10 minutes on high with 30 seconds on and 30 seconds off, 3 times. The samples were clarified by centrifugation at 13,000 × g for 15 minutes at 4°C. Following protein quantification, 600 μg of lysate was incubated with 5 μg of either polyclonal rabbit anti-SBDS (Abcam, ab128946) or polyclonal rabbit anti-RPS19 (Bethyl-A304-002A) antibody overnight at 4°C with rotation. An aliquot was incubated with IgG (Cell Signaling, cat# 2729) as a control to determine background binding. Pierce protein G magnetic beads (Thermo Fisher Scientific, cat# 88848) were added and the samples were rotated for 1 hour at 4°C. The beads were washed twice in RIPA buffer, 4 times in wash buffer (100 mM Tris-HCl at pH 8.5, 500 mM LiCl, 1% NP-40, 1% sodium deoxycholate) followed by 2 washes in RIPA buffer and twice with 1X Tris-EDTA (TE). Immunocomplexes were eluted twice by adding 200 μL of elution buffer (1X TE, 1% SDS) and incubating for 15 minutes at 65°C. Cross-links were reversed by incubation for 7 hours at 65°C, with a final concentration of 200 mM NaCl. The samples were then treated with RNase A (20 μg) and digested with proteinase K (40 μg). DNA was purified with phenol/chloroform precipitation and used as a template for real-time PCR. The primer pairs used to analyze the −2 Kb, −1 Kb and +5 kb regions and GAPDH were as previously published (46): −2 Kb [forward (for)] 5’-GCCCATATATGGAGTTCCGC-3’ and [reverse (rev)] 5’- CGTAAGGTCATGTACTGGGC-3’; −1 Kb (for) 5’-GACCGCGTTACTCCCACAGGTG-3’ and (rev) 5’-GGCTTTCACGCAGCCACAGAAAA3’; +5 Kb (for) 5’- CACTAACTTGCTCACACTATCCTCG-3’and (rev) 5’- TTTCTAGACCAGCCCACGTAATG; and GAPDH (for) 5’ CACATGGCCTCCAAGGAGTAAG-3’ and (rev) 5’-TACATGACAAGGTGCGGCTCCC- 3’. Primers used to analyze the +1 Kb region were as previously published (47) (for) 5’- TGTGGTTTCCAAATGTGTCAG-3’ and (rev) 5’-TACCCGCTTCCATTGCTC-3’.

### Olaparib sensitivity assays

Twenty-four hours following siRNA transfections as described previously, U2OS cells were re-plated in 96 well plate wells at a density of 2000 cells per well. Olaparib (Selleck Chemical) was added to final concentrations of 0.75, 1.5, 3 and 6 μM or DMSO alone and the cells were incubated for 48 hours. Cell viability was assessed using CellTiter 96® Aqueous Non-Radioactive Cell Proliferation reagent (Promega, G5421).

### Statistics

The statistical tests are indicated in the respective figure legends. Error bars indicate mean ± standard deviation (SD). *P* values of 0.05 or less were considered significant. Analyses were performed using GraphPad Prism (GraphPad Software). For microscopy data, the 1-way analysis of variance (ANOVA) with Dunnett’s multiple comparisons test was used. For flow cytometry data, 1-way ANOVA and Dunnett’s or Tukey’s multiple comparisons test was used. For cell cycle analysis, 2-way ANOVA and Dunnett’s multiple comparisons was used. For western blot analysis, 1-way ANOVA with Dunnett’s multiple comparisons test was used. For ChIP analysis, t test was used.

### Study approval

This investigation was conducted according to the Declaration of Helsinki Principles. Written informed consent was received from participants before inclusion in the study according to protocol H-17698 Genetic and Biological Determinants of Bone Marrow Failure approved by the Institutional Review Board for Baylor College of Medicine and affiliated hospitals.

## AUTHOR CONTRIBUTIONS

A.A.B. designed the research studies, analyzed the data, and co-wrote the manuscript.

E.A. designed the research studies, performed all of the experiments except the colony survival assays, analyzed the data and co-wrote the manuscript.

N.C. performed the colony survival assays and analyzed the data.

## ACKNOWLEDGMENTS

We thank the research subjects for their participation and the clinical research staff for their assistance. We also thank members of the Bertuch lab for helpful discussions throughout this project and critical reading of the manuscript. This study was funded by grants to A.A.B. from the Shwachman-Diamond Syndrome Foundation and Hyundai Hope on Wheels. Lymphoblastoid cell line generation was supported in part by the Clinical Translational Core of the Baylor College of Medicine Intellectual and Developmental Disabilities Center (P50HD103555) from the Eunice Kennedy Shriver National Institute of Child Health and Human Development. Imaging for this project was supported by the Integrated Microscopy Core at Baylor College of Medicine and the Center for Advanced Microscopy and Image Informatics (CAMII) with funding from NIH (DK56338, CA125123, ES030285), and CPRIT (RP150578, RP170719), the Dan L. Duncan Comprehensive Cancer Center, and the John S. Dunn Gulf Coast Consortium for Chemical Genomics.

