## Supplemental Tables and Figure for "Homologous recombination defects in Shwachman-Diamond syndrome and Diamond-Blackfan anemia"

**Supplemental Table 1 SDS and DBA LCLs used in this study**

| **LCL ID** | **Mutation(s)** |
|---|---|
| **SDS LCLs** | |
| BMF30 | *SBDS* c.183_184 TA>CT; c.258+2 T->C |
| BMF32 | *SBDS* c. p.K33E/wt and structural variation |
| BMF40 | *SBDS* c.183_184 TA>CT; c.258+2 T->C |
| BMF94 | *SBDS* c.183_184 TA>CT; c.258+2 T->C |
| BMF113 | *SBDS* c.183_184 TA>CT; c.258+2 T->C |
| **DBA LCLs** | |
| BMF16 | *RPL5* c.535C>T; p.R179* |
| BMF51 | *RPS19* c.1-1G>A |
| BMF54 | *RPS19* c.280C>T; p.R94* |
| BMF74 | *RPS26* c.1A>G; p.M1V |
| BMF78 | Clinical diagnosis DBA, genetically uncharacterized |
| BMF89 | *RPL5* c.70C>T; p.R24* |
| BMF92 | *RPS20* c.251A>T; p.Ile84Asn |

**Supplemental Table 2 SiRNAs used in this study**

| siRNA target[1] | Sequence | Sequence | Catalog number |
|---|---|---|---|
| Universal Negative Control #1 | | | SIC001 |
| RAD51 | AGAAGGAGCUAAUAAAUAUtt | AUAUUUAUUAGCUCCUUCUtt | s531930 |
| SBDS-1 | CAUACACCGUGAUCCUUAUtt | AUAAGGAUCACGGUGUAUGgt | s27484 |
| SBDS-2 | GGUUCUUUGGAAGUACUCAtt | UGAGUACUUCCAAAGAACCtt | s27483 |
| EFTUD1 | GGAAGUACAUGAACGCAGUtt | ACUGCGUUCAUGUACUUCCgg | s35960 |
| EFTUD2 | CCAACAUACUAGUCAAUAAtt | UUAUUGACUAGUAUGUUGGgc | s35959 |
| RPS19-1 | GAUGAGAACUGGUUCUACAtt | UGUAGAACCAGUUCUCAUCgt | s535016 |
| RPS19-2 | GCCGCAAACUGACACCUCAtt | UGAGGUGUCAGUUUGCGGCcg | s12323 |
| eIF6 | CCUACUGUCUGGUAGCGAUtt | AUCGCUACCAGACAGUAGGtg | s7586 |

[1]All siRNAs obtained from Thermo Fisher (Ambion, Silencer select) except for the Universal Negative Control #1, which was obtained from Millipore Sigma.

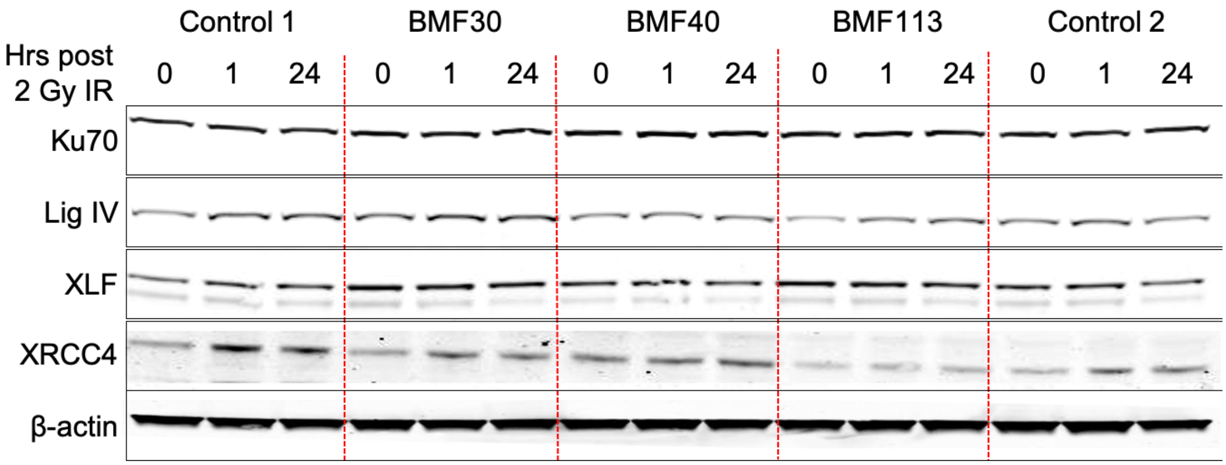

**Supplemental Figure 1 NHEJ protein levels in SDS LCLs are similar to normal control LCLs** Representative western blot of WCEs from 3 SDS (BMF113, BMF30 and BMF40) and 2 unique control LCLs prepared pre-treatment (0) and following treatment with 2 Gy IR harvested at 1 and 24 hours analyzed for the designated NHEJ. β-actin is a loading control.
